# Fgf signalling is required for gill slit formation in the skate, *Leucoraja erinacea*

**DOI:** 10.1101/2023.08.20.554054

**Authors:** Jenaid M. Rees, Michael A. Palmer, J. Andrew Gillis

## Abstract

The gill slits of fishes develop from an iterative series of pharyngeal endodermal pouches that contact and fuse with surface ectoderm on either side of the embryonic head. We find in the skate (*Leucoraja erinacea*) that all gill slits form via a stereotypical sequence of epithelial interactions: **1)** endodermal pouches approach overlying surface ectoderm, with **2)** focal degradation of ectodermal basement membranes preceding endoderm–ectoderm contact; **3)** endodermal pouches contact and intercalate with overlying surface ectoderm, and finally **4)** perforation of a gill slit occurs by epithelial remodelling, without programmed cell death, at the site of endoderm– ectoderm intercalation. Skate embryos express *Fgf8* and *Fgf3* within developing pharyngeal epithelia during gill slit formation. When we inhibit Fgf signalling by treating skate embryos with the Fgf receptor inhibitor SU5402 we find that endodermal pouch formation, basement membrane degradation and endodermal–ectodermal intercalation are unaffected, but that epithelial remodelling and gill slit perforation fail to occur. These findings point to a role for Fgf signalling in epithelial remodelling during gill slit formation in the skate and, more broadly, to an ancestral role for Fgf signalling during pharyngeal pouch epithelial morphogenesis in vertebrate embryos.

## Introduction

The pharyngeal arches of vertebrates are a series of paired columns of tissue that form on either side of the embryonic head. Pharyngeal arches are delineated by an iterative series of endodermal pouches that outpocket laterally from the foregut endoderm and that contact overlying surface ectoderm (Fig. 1A,B). The precise number of pharyngeal pouches and arches varies across vertebrate taxa (Graham et al., 2019), but pouches generally form in an anterior-to-posterior sequence, and resulting arches are lined laterally by ectoderm, medially by endoderm, and contain a core of mesoderm and mesenchyme, the latter of both neural crest and lateral mesodermal origin (Graham and Smith, 2001, Sleight and Gillis, 2020). In all vertebrates, pharyngeal arch ectoderm produces epidermis and sensory neurons, while the pharyngeal arch core mesoderm and mesenchyme give rise to musculature, endothelia, and skeletal/connective tissues of the head (Graham and Smith, 2001). The fate of pharyngeal arch endodermal pouches, on the other hand, differs between fishes and amniotes: in fishes, endodermal pouches fuse with the overlying surface ectoderm, giving rise to the gill slits and the respiratory lamellae of the gills (Gillis and Tidswell, 2017 - Fig. 1C), while in amniotes, the endodermal pouches give rise to glandular tissues, such as the tonsils, and parathyroid and ultimobranchial glands (Grevellec and Tucker, 2010).

**Figure 1:**
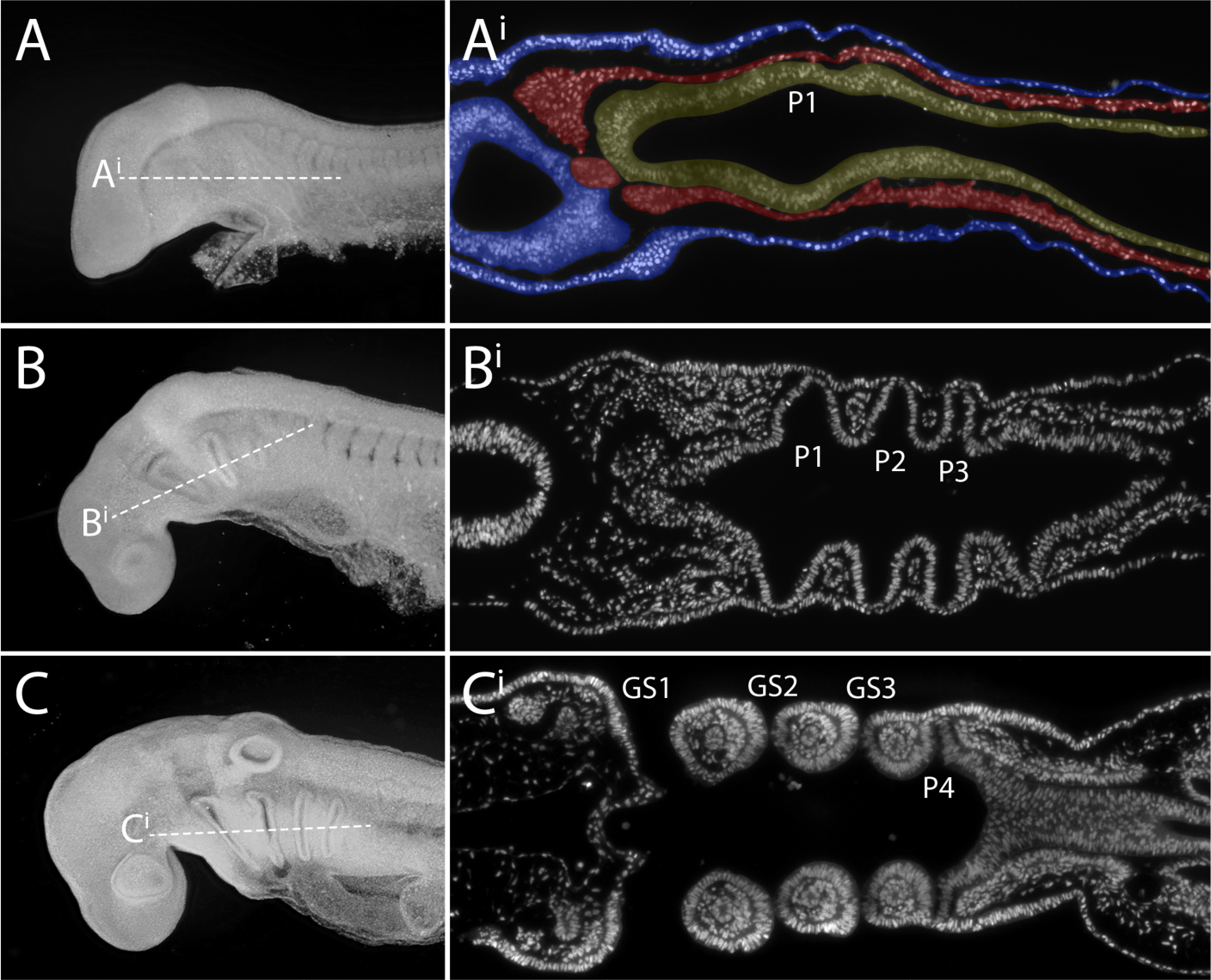
Overview of pharyngeal arch development in the little skate. Skate embryos at stages (S)18, 20 and 23 in lateral view (anterior to the left), and corresponding horizontal sections through the pharynx, stained with DAPI. **(A–Ai)** The first pharyngeal pouch begins to form as an outpocket from the pharyngeal endoderm at S18. Additional pouches will follow in an anterior-to-posterior sequence. **(B–Bi)** By S20, three pharyngeal pouches have formed, and these pouches are in contact with surface ectoderm. **(C–Ci)** By S23, the first, second and third pharyngeal pouches have fused with surface ectoderm to form gill slits, while the fourth pouch has contacted the surface ectoderm, but has not yet fused. In **Ai**, endoderm is false-coloured yellow, mesoderm is false-coloured red, and ectoderm is false-coloured blue. *GS1-GS3*, gill slits 1-3; *P1-P4*, pharyngeal pouches 1-4. Scale bars: A,B,C = 500µm; Ai,Bi,Ci = 100µm.

In fishes, Fgf signalling is a key regulator of pharyngeal endodermal pouch development. Crump et al. (2004) found that a reduction in Fgf signalling in zebrafish – either by *fgf3* morpholino knock-down in an *fgf8* null mutant background, or by treatment of embryos with the small molecule Fgf receptor inhibitor SU5402 – results in a complete loss of pharyngeal endodermal pouches, and that restoration of wildtype *fgf* expression in mesoderm and neural tissue by blastomere transplantation rescues this pharyngeal pouch defect. Zebrafish endodermal pouches undergo lateral migration following destabilizing non-canonical Wnt11r-mediated signalling from adjacent pharyngeal mesoderm (Choe et al., 2013), with mesodermal Fgf8 signalling functioning as a chemoattractant guidance cue to direct this lateral migration of pouch endoderm (Choe and Crump, 2014). Pharyngeal mesodermal expression of *wnt11r* and *fgf8* are both downstream of mesodermal Tbx1, corroborating and providing mechanistic insight into pharyngeal segmentation defects that were previously observed in *tbx1*^-/-^ fish (Piotrowski and Nusslein-Volhard, 2000; Piotrowski et al., 2003). Fgf signalling is similarly required for pharyngeal pouch formation in jawless fishes. In the lamprey, *fgf3* and *fgf8* are expressed in pharyngeal epithelia, and inhibition of FGF signalling in lamprey embryos results in a loss of pharyngeal pouch formation (Jandzik et al., 2014).

Fgf8 signalling from pharyngeal epithelia is also implicated in pharyngeal pouch development in mammals. In *Foxi3*^-/-^ mice, there is a delay in onset of *Fgf8* expression in pharyngeal endoderm and ectoderm, and this leads to profound craniofacial skeletal mispatterning (Edlund *et al*., 2014). But unlike in zebrafish, where pharyngeal pouches fail to form in the absence of Fgf signalling, formation of the first pharyngeal pouch in mouse appears to be laregly robust to reductions in Fgf8 dosage. Instead, Fgf8 signalling is required in mouse for separation of the mandibular and hyoid arches via reshaping and elongation of the first pharyngeal pouch and interaction of this pouch with adjacent ectoderm (Zbasnik, 2023).

While FGF signalling clearly played an important role in pharyngeal endodermal pouch formation and/or morphogenesis in the last common ancestor of vertebrates, the molecular control of ultimate fusion of endodermal pouches with surface ectoderm and perforation of gill slits in fishes remains unexplored. A comparative analysis of endodermal pouch development in chick and shark embryos identified displacement of the ectoderm by outgrowing endodermal pouches as a conserved feature of endoderm–ectoderm fusion in the pharynx of jawed vertebrates (Shone and Graham, 2014). In the chick, there is a single instance of fusion at the level of the 2^nd^ pharyngeal pouch, with the endoderm and ectoderm first contacting via cellular processes. These processes interdigitate to form a single epithelial layer that thins and perforates, with perforations enlarging until a connection between the pharynx and the external environment has formed either by epithelial remodelling (Waterman, 1985) or by cell death (Shone and Graham, 2014). In the shark, on the other hand, it has been reported that each endodermal pouch pushes through the surface ectoderm before perforating to form gill slits (Shone and Graham, 2014). How epithelial events leading to pharyngeal endoderm–ectoderm fusion and gill slit formation are influenced by signals from adjacent tissues is not known.

Here, we investigate the epithelial and signalling interactions that form the basis of gill slit formation in embryos of a cartilaginous fish, the little skate (*Leucoraja erinacea*). We find in the skate that as pharyngeal endodermal pouches approach overlying surface ectoderm, focal degradation of the ectodermal basement membrane precedes and predicts impending points of endoderm–ectoderm contact. Cytoskeletal protrusions then extend from the ectoderm toward pouch endoderm, and this is followed by endodermal basement membrane degradation, and by ectoderm– endoderm contact and intercalation. Gill slit perforation then occurs within the intercalated endoderm–ectoderm by epithelial remodeling without programmed cell death, closely resembling the process described at the level of the 2^nd^ pharyngeal pouch of chick embryos by Waterman (1985). We find that genes encoding Fgf ligands are expressed in pharyngeal endoderm and ectoderm in skate, and by pharmacologically inhibiting Fgf signalling we find that this pathway is dispensable for endoderm–ectoderm intercalation but is required for subsequent epithelial remodelling and gill slit perforation. These findings provide the first insight into a molecular mechanism and tissue interactions controlling gill slit formation in a jawed vertebrate and expand the roles of FGF signalling during early pharyngeal arch development in vertebrates.

## Methods

### Skate embryos and pharmacological manipulations

Little skate (*Leucoraja erinacea)* eggs were obtained from the Marine Resources Center at the Marine Biological Laboratory in Woods Hole, MA, U.S.A. Eggs were reared in artificial seawater at 15°C and staged according to Ballard *et al*. (1993). For treatment of skate embryos with SU5402 (Sigma Aldrich), skate egg cases containing embryos at stage (S)19 or S20 were injected with 20µL of 25mM SU5402 in dimethyl sulfoxide (DMSO) or 20µL of DMSO alone (as control) using a 1mL syringe and 30-gauge needle. Skate egg cases hold a mean volume of ∼10mL, resulting in a final *in ovo* concentration of 50μM SU5402. Eggs were gently rocked to distribute the drug or control vehicle throughout the egg, and eggs were then reared for 4 days (as survival rapidly decreased beyond this point). After 4 days, embryos were euthanised with an overdose of MS-222 (1g/L in seawater) and fixed in 4% paraformaldehyde (Electron microscopy sciences) as per Gillis *et al*. (2012). SU5402 treatments were attempted using this same strategy at S18, but embryos treated at this stage did not survive beyond 48 hours post-injection.

### Paraffin embedding, sectioning, and histochemical staining

Embryos were embedded in paraffin and sectioned at 7μm as described in O’Neill *et al*.(2007). Sectioned embryos were stained with Hematoxylin and Eosin by dewaxing in two rinses of Histosol (National Diagnostics), rehydration through serial dilutions of ethanol, then staining for 15 minutes in Mayer’s Haematoxylin (Sigma Aldrich). Slides were rinsed in running tap water for 20 minutes, washed briefly in 95% ethanol and stained with 0.1% w/v Eosin Y (Sigma Aldrich) in 95% ethanol for 2 minutes. Slides were then rinsed in 100% ethanol, cleared with Histosol and mounted with DPX (Sigma Aldrich).

### Gelatin embedding, sectioning, and immunofluorescent staining

Embryos were embedded in 15% w/v gelatin and sectioned at 220μm using a vibratome. These sections were washed and permeabilized in PBS + 0.1% triton (PBST), and then blocked in 1% bovine serum albumin in PBST before addition of primary antibodies against E-cadherin (1:200, BD Transduction Laboratories, 610181), laminin (1:200, Sigma-Aldrich, L9393), and/or cleaved caspase-3 (1:200, Cell Signaling Technology, D175). Sections were incubated in primary antibody overnight at 4°C. The primary antibodies were then washed off with PBST and then sections were incubated in secondary antibodies (1:200, AlexaFluor-conjugated, ThermoFisher) and phalloidin (1:400, ThermoFisher, A12379) overnight at 4°C. Finally, secondary antibodies were washed off with PBST and sections were then graded into 75% glycerol for imaging.

### mRNA *in situ* hybridisation

Chromogenic mRNA *in situ* hybridisation (ISH) was performed on paraffin sections and in wholemount as described in O’Neill *et al*. (2007), with modifications according to Gillis *et al*. (2012). ISH probes were generated against skate *Fgf8* (GenBank EU574737.1) and *Fgf3* (GenBank XM_055649200) by *in vitro* transcription using standard methods as described by O’Neill *et al*. (2007). Third-generation mRNA ISH by chain reaction (HCR) was performed following the Choi *et al*. (2018) protocol for formaldehyde fixed, paraffin-embedded sections, with modifications as per Criswell and Gillis (2020). Probe sets for *Dusp6* (Lot PRA756) and *Fgf8* (Lot PRA755), buffers, and hairpins were purchased from Molecular Instruments (Los Angeles, California, USA). Following ISH, slides were rinsed in PBS, post-fixed in 4% paraformaldehyde and coverslipped with DAPI-Fluoromount® G (SouthernBiotech), while wholemounts were rinsed, post-fixed and graded into 75% glycerol for imaging.

### TUNEL Staining

Embryos were paraffin embedded and sectioned as described above, and TUNEL staining was performed using the Fluorescein *in situ* Cell Death Detection Kit (Sigma-Aldrich) according to the manufacturer’s instructions. Sections were coverslipped with Fluoromount-G containing DAPI prior to imaging. Sections at anatomically equivalent mid-points in the posterior pharyngeal arches were selected and imaged.

### Imaging

Paraffin embedded tissue sections were imaged on a Zeiss Axioscope.A1 compound microscope with a Zeiss colibri 7 fluorescence LED light source using a Zeiss Axiocam 305 colour or 503 mono camera and ZenPro software. Gelatin embedded tissue was imaged on Zeiss LSM 780 confocal laser scanning microscope. Whole embryos were imaged using a Leica M165FC stereomicroscope, a Leica DFC7000 T camera and LAS X software. All figures were assembled using Fiji and Adobe creative cloud. Images were flipped for consistency where needed.

## Results

### Epithelial morphogenesis during gill slit formation in the skate

To image the interactions of pharyngeal endoderm and ectoderm during gill slit formation in the skate, we sectioned and stained skate embryos from stages (S)18– S22. These stages span early development of the pharyngeal arches, from the onset of endodermal pouch formation to presence of perforated gill slits. We first co-stained sections for e-cadherin and phalloidin to visualise cell–cell junctions and filamentous actin, respectively, and to obtain a general overview of endoderm–ectoderm interactions during gill slit formation. We found that as pouches bulge out laterally from the pharyngeal endoderm (Fig. 2A), they displace the mesoderm layer that sits between the endoderm and ectoderm (Fig. 2B), and eventually contact the overlying surface ectoderm (Fig. 2C). Pouch endoderm and ectoderm, once in close apposition, fuse into a single disorganised epithelium (Fig. 2D). This epithelium then perforates (Fig. 2E), giving rise to a complete gill slit that opens from the pharyngeal cavity to the external environment (Fig. 2F). The images shown in Fig. 2 are representative of observations made at the level of each pharyngeal pouch, and the tissue interactions described above appear to be general for all developing gill slits.

**Figure 2.**
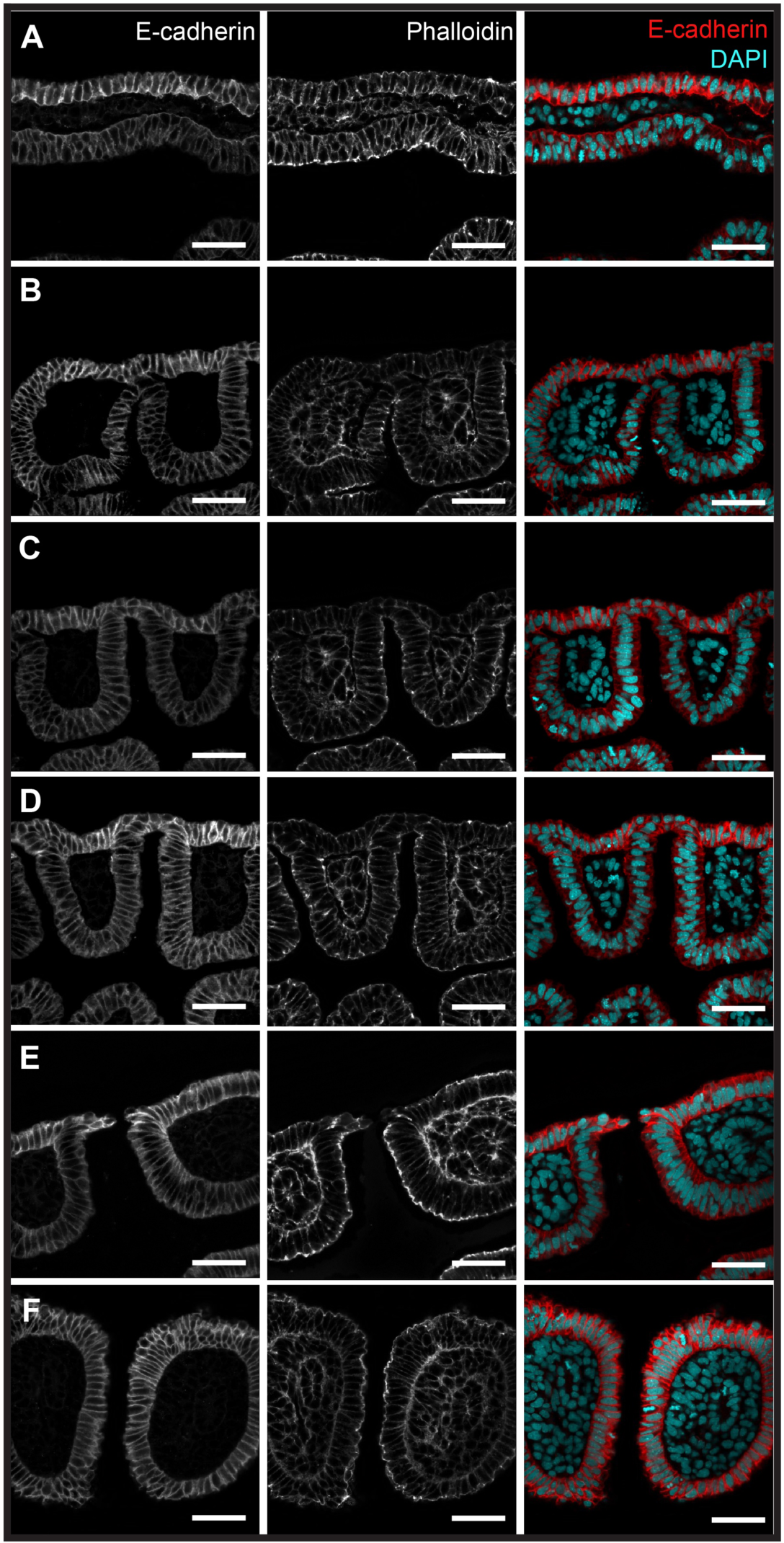
Sequence of epithelial interactions during gill slit development in the little skate. Immunofluorescence for E-cadherin and phalloidin staining of developing pharyngeal pouches and gill slits in the skate. **(A)** Endodermal pouches initially bulge out toward the overlying surface ectoderm, displacing the mesoderm layer that sits between the endoderm and ectoderm. **(B)** The endoderm then contacts the overlying ectoderm and **(C)** begins to push against the surface ectoderm. **(D)** Endoderm and ectoderm form a single epithelium. **(E)** The epithelium perforates at the site of endoderm-ectoderm contact, forming a gill slit opening. **(F)** The columns of tissue that are delineated by adjacent gill slits are called pharyngeal arches. All scale bars = 50 µm.

Prior to fusion of the endoderm and ectoderm, both tissue layers possess intact basement membranes. Using laminin immunofluorescence, we followed the integrity of these basement membranes. We found that prior to endoderm–ectoderm contact, degradation occurs first in the basement membrane of the ectoderm, with ectodermal cytoskeletal protrusions extending through newly formed holes in the basement membrane toward the endoderm (Fig. 3A). These protrusions contact the endoderm and form bridges between the two tissue layers. As more bridges form, the basement membrane of the endoderm degrades, and this permits intercalation of cells from ectodermal and endodermal epithelia to begin (Fig. 3B). The resulting intercalated epithelium extends between two adjacent pharyngeal arches. Prior to gill slit perforation, the endodermal and ectodermal basement membranes on either side of the intercalated epithelium join up, resulting in an intact basement membrane underlying the epithelium of each pharyngeal arch (Fig. 3B,C). The complete basement membrane within the pharyngeal arch remains intact throughout the process of perforation and persists in the final arch structure (Fig. 3D)

**Figure 3.**
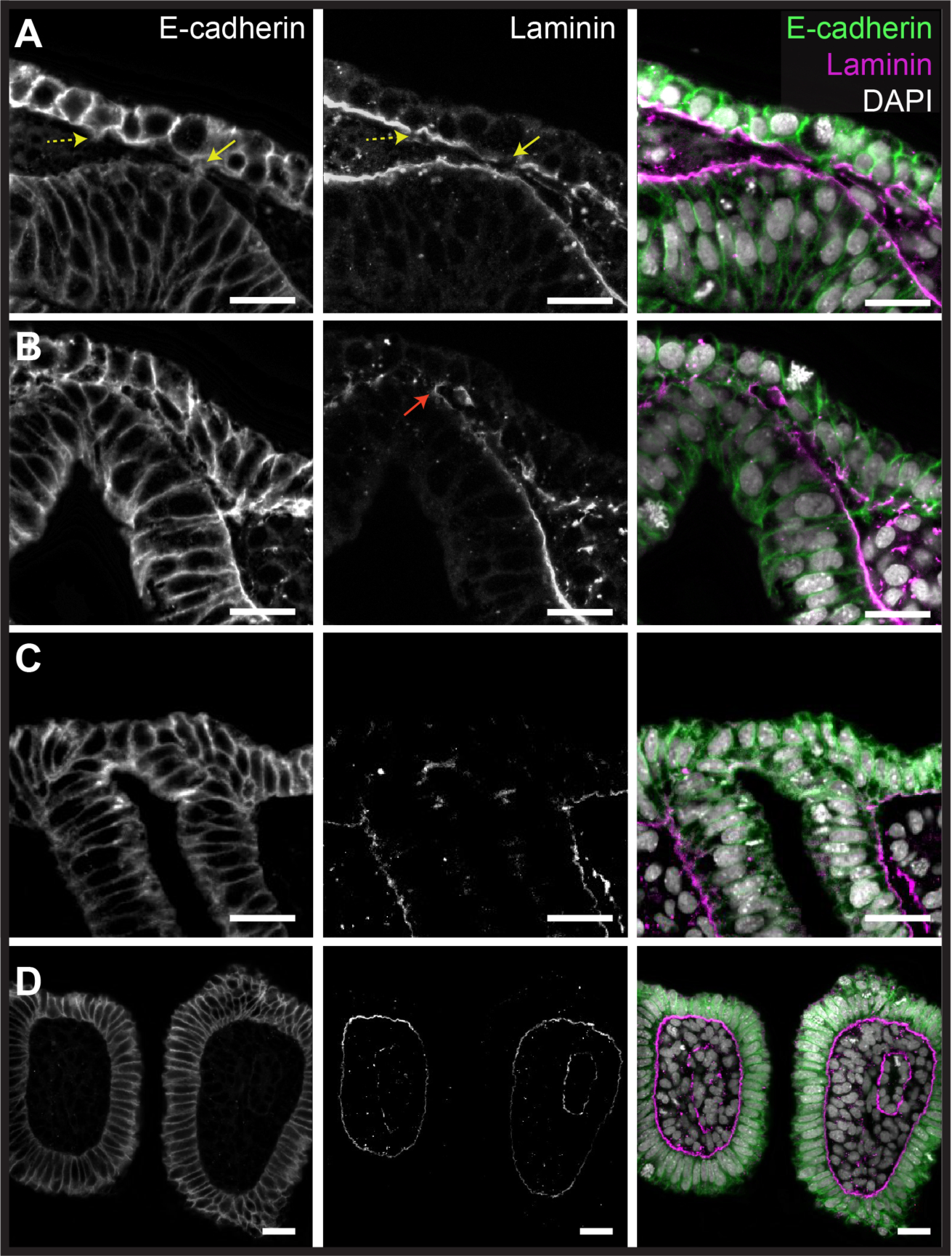
Basement membrane degradation and epithelial protrusions precede endoderm-ectoderm intercalation and gill slit formation. Immunofluorescence for E-cadherin and laminin reveals that **(A)** degradation of the ectodermal basement membrane precedes endoderm-ectoderm contact, with small cytoskeletal protrusions (dashed yellow arrow) forming bridges (yellow arrow) between the ectoderm and the endodermal basement membrane. **(B)** These bridges continue to form and are associated with further basement membrane degradation in the ectoderm and endoderm. The endodermal and ectodermal basement membranes flanking the point of endoderm-ectoderm contact join to form a new continuous basement membrane (red arrow) **(C)** The basement membrane fully degrades at the site of endoderm-ectoderm contact, leading to intercalation of the endoderm and ectoderm. At this stage, endoderm and ectoderm are no longer recognisable as distinct epithelia. **(D)** After perforation of the gill slit is complete, the new continuous endodermal-ectodermal basement membrane flanking the gill slit forms persists within the pharyngeal arch. All scale bar = 25 µm.

### Skate gill slit perforation occurs by epithelial remodelling without cell death

Shone and Graham (2014) reported that foci of cell death appear prior to ectodermal displacement at the level of the 2^nd^ and 3^rd^ pharyngeal pouches in the chick, based on retention of the vital dye Lysotracker Red. To determine whether programmed cell death is a general mechanism of pharyngeal epithelial perforation for jawed vertebrates, we used TUNEL staining and immunofluorescence for cleaved caspase-3 to test for apoptosis in conjunction with epithelial remodelling during gill slit development in the skate. We conducted these analyses at S22 because this stage includes pharyngeal pouches at all phases of development/epithelial interaction (i.e., pre-endoderm–ectoderm contact, during endoderm–ectoderm contact and intercalation, and recently perforated gill slits). TUNEL staining was performed on paraffin sections, and all sections through the pharynx were analysed for each embryo (n=4 embryos), while immunofluorescence for cleaved caspase-3 was performed on vibratome sections and imaged by confocal microscopy (n=3 embryos). While positive TUNEL and cleaved caspase-3 staining were observed in nearby rostral pharyngeal epithelium at S22 (Fig. 4A, Bi,Bii; Fig. S1A), we observed no positive TUNEL or cleaved caspase-3 staining during endoderm–ectoderm contact, epithelial intercalation or gill slit perforation (Fig. 4C–Dii; Fig. S1B,C). This indicates that unlike reports from chick (Shone and Graham, 2014), cell death does not occur during gill slit formation in skate embryos, and that gill slit perforation instead occurs by epithelial remodelling without cell death.

**Figure 4:**
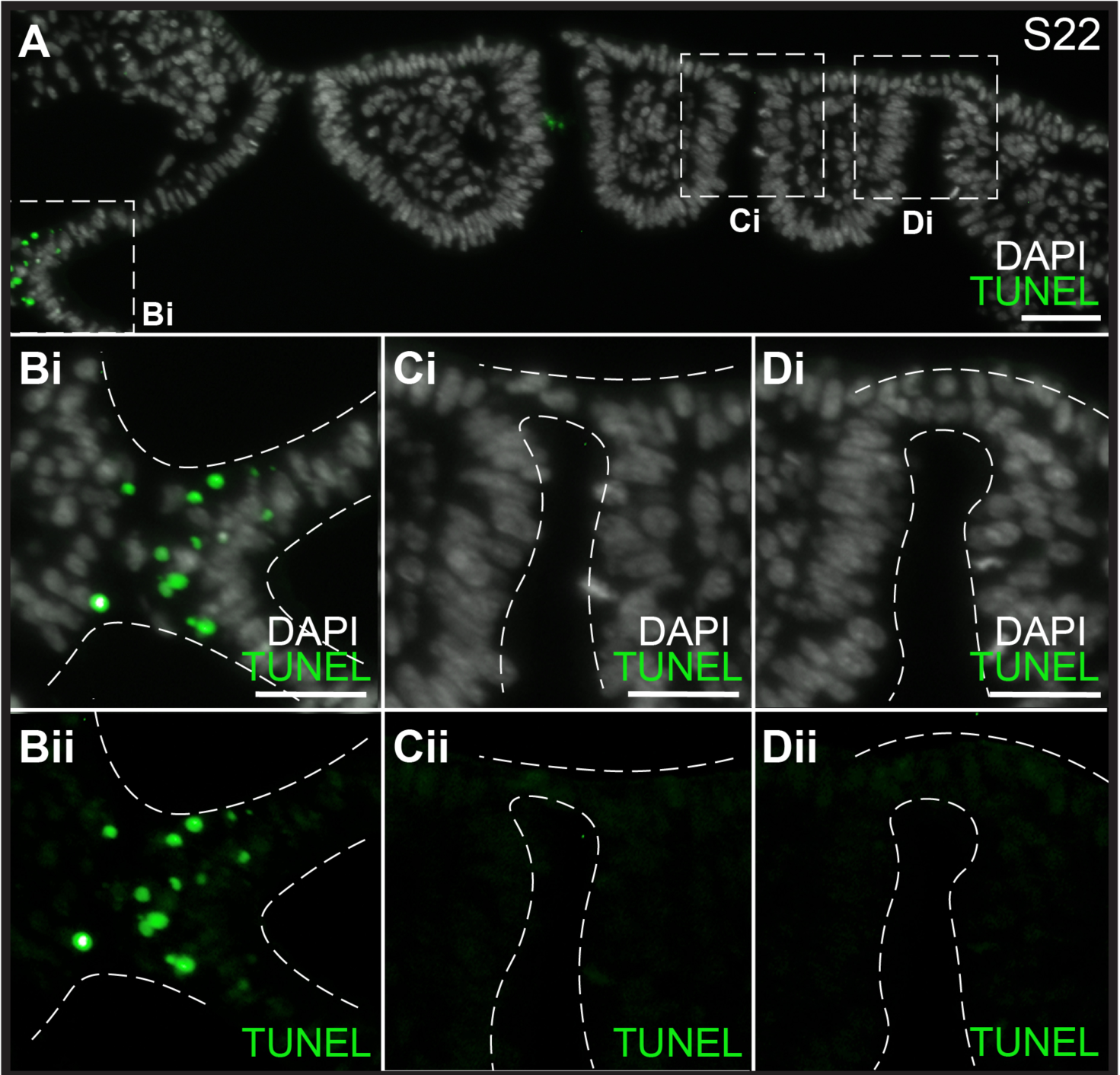
Cell death does not contribute to gill slit perforation in the skate. **(A)** Horizontal section through a S22 skate embryo stained with TUNEL and DAPI. (**Bi-Bii**) There is a clear region of positive TUNEL staining indicating cell death between the rostral pharyngeal epithelium and the neural tube. However, (**Ci/Cii-Di/Dii**) no cell death was detected at any stage during endoderm–ectoderm intercalation or gill slit perforation. All sections were stained and imaged for n=4 S22 skate embryos. Scale bars: A = 50µm, Bi-Di = 25µm.

### Expression of *Fgf8* and *Fgf3* during gill slit development in the skate

Given established roles for Fgf8 and Fgf3 signalling in the control of pharyngeal pouch development or morphogenesis in zebrafish (Crump et al., 2004), lamprey (Jandzik et al., 2014) and mouse (Zbasnik, 2023), we examined spatial expression of *Fgf8* and *Fgf3* by mRNA *in situ* hybridisation during skate gill slit development. We found that discrete *Fgf8* expression domains appear in a rostral-to-caudal sequence, in the vicinity of each developing pharyngeal pouch, from S18–S22 (Fig. 5A–E). This expression is first detected in the caudal endoderm of each developing pharyngeal pouch (Fig. 5AF), and later also in the overlying surface ectoderm at the level of pouches that are intercalating with surface ectoderm (Fig. 5G). We observed similar rostral-to-caudal appearance of discrete *Fgf3* expression domains associated with each developing pharyngeal pouch from S18–S22 (Fig. 5H–L), though unlike *Fgf8* expression, *Fgf3* was expressed only in the caudal endoderm of each pharyngeal pouch (Fig. 5M,N). There is a distinct boundary between the *Fgf8*- and *Fgf3*-negative anterior pouch epithelium and *Fgf8*- and *Fgf3*-positive posterior pouch epithelium, and this boundary corresponded with the site of future gill slit perforation. At no point from S18–S22 did we detect expression of *Fgf8* or *Fgf3* in pharyngeal mesoderm or mesenchyme. Collectively, these expression patterns indicate that Fgf signalling may be involved in aspects of pharyngeal pouch or gill slit development in the skate, with skate *Fgf8* and *Fgf3* expression most closely resembling the pharyngeal epithelial expression patterns previously reported in lamprey.

**Figure 5.**
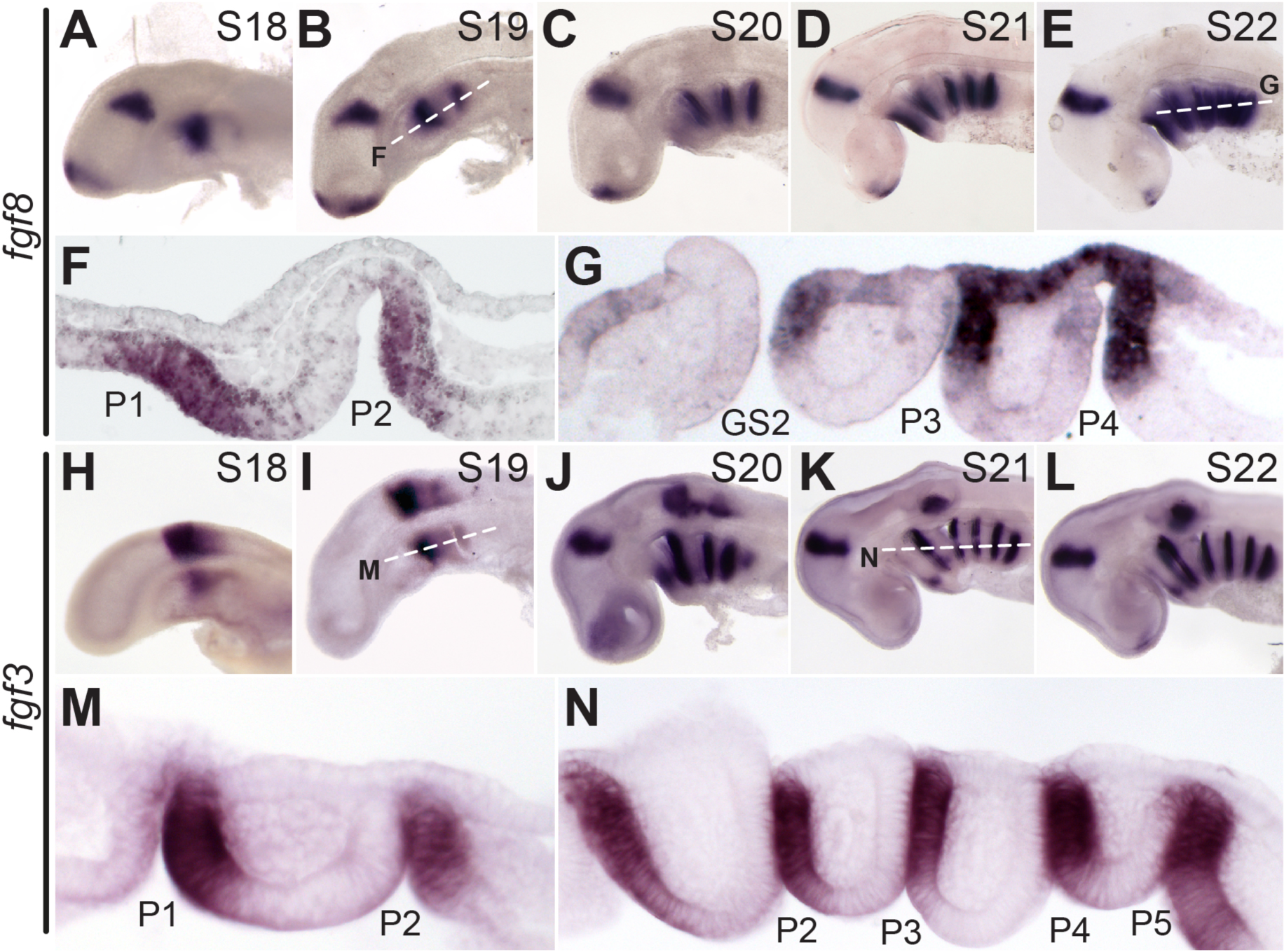
*Fgf8* and *Fgf3* are expressed in the skate endoderm and ectoderm during pharyngeal pouch formation and fusion. **(A–E)** *Fgf8* is expressed sequentially with the formation of pharyngeal endodermal pouches, as shown by wholemount mRNA ISH at S18–22. Horizontal sections of **(F)** S19 and **(G)** S22 embryos indicate that *Fgf8* is expressed in the posterior endodermal epithelium of each pharyngeal pouch and in the overlying ectodermal epithelium during endoderm-ectoderm intercalation. **(H–L)** *Fgf3* is expressed sequentially with the formation of pharyngeal endodermal pouches, as shown by wholemount mRNA ISH at S18–22. Horizontal sections of **(M)** S19 and **(N)** S22 embryos indicate that *Fgf3* is expressed in the posterior endodermal epithelium of each developing pharyngeal pouch. *GS2*, gill slit 2; *P1-P5*, pharyngeal pouches 1–5. Scale bars; A–E and H-L = 500µm; F,G,M,N = 100µm.

### Fgf signalling is required for gill slit perforation in skate

Finally, to test whether Fgf signalling is required for gill slit development in skate, we inhibited Fgf signalling during early pharyngeal arch development by *in ovo* delivery of SU5402, a small molecule inhibitor of Fgf receptor tyrosine kinase activity (Mohammadi et al., 1997). To confirm that our *in ovo* treatment method effectively inhibits FGF signalling in the skate, we tested for expression of *Dusp6* 24 hours post-administration of SU5402 or control and noted a substantial reduction in *Dusp6* transcription with SU5402 treatment (Fig. S2). We were able to treat embryos *in ovo* from S19/20 and achieve high rates of survival for 4 days post injection, to S22. At S19/20, skate embryos possess two or three pharyngeal pouches (Fig. S3), but importantly, there are no perforated gill slits at these stages.

Embryos treated with SU5402 from S19/20 showed no significant difference in mean number of pharyngeal pouches relative to controls at S22 (n=15 SU5402-treated; n=16 control; 3.3 vs 3.75 mean pouches; p=0.06; t-test) (Fig. 6A), suggesting that SU5402 treatment does not cause a general delay in pharyngeal development. However, we found that SU5402-treatment from S19/20 resulted in a significant reduction in the number of gill slits at S22 (n=15 SU5402-treated; n=16 control; 0.06 vs 1.75 mean gill slits; p=2.21e^-12^; t-test) (Fig. 6B). Of all embryos treated with SU5402, only one embryo showed perforation of the 1^st^ gill slit, while no SU5402-treated embryos showed perforation of the 2^nd^ gill slit. This contrasted with control embryos, where the 1^st^ gill slit was perforated in all embryos, and the 2^nd^ gill slit was perforated in 80% of embryos (n=12/16) (Fig. 6C,D).

**Figure 6.**
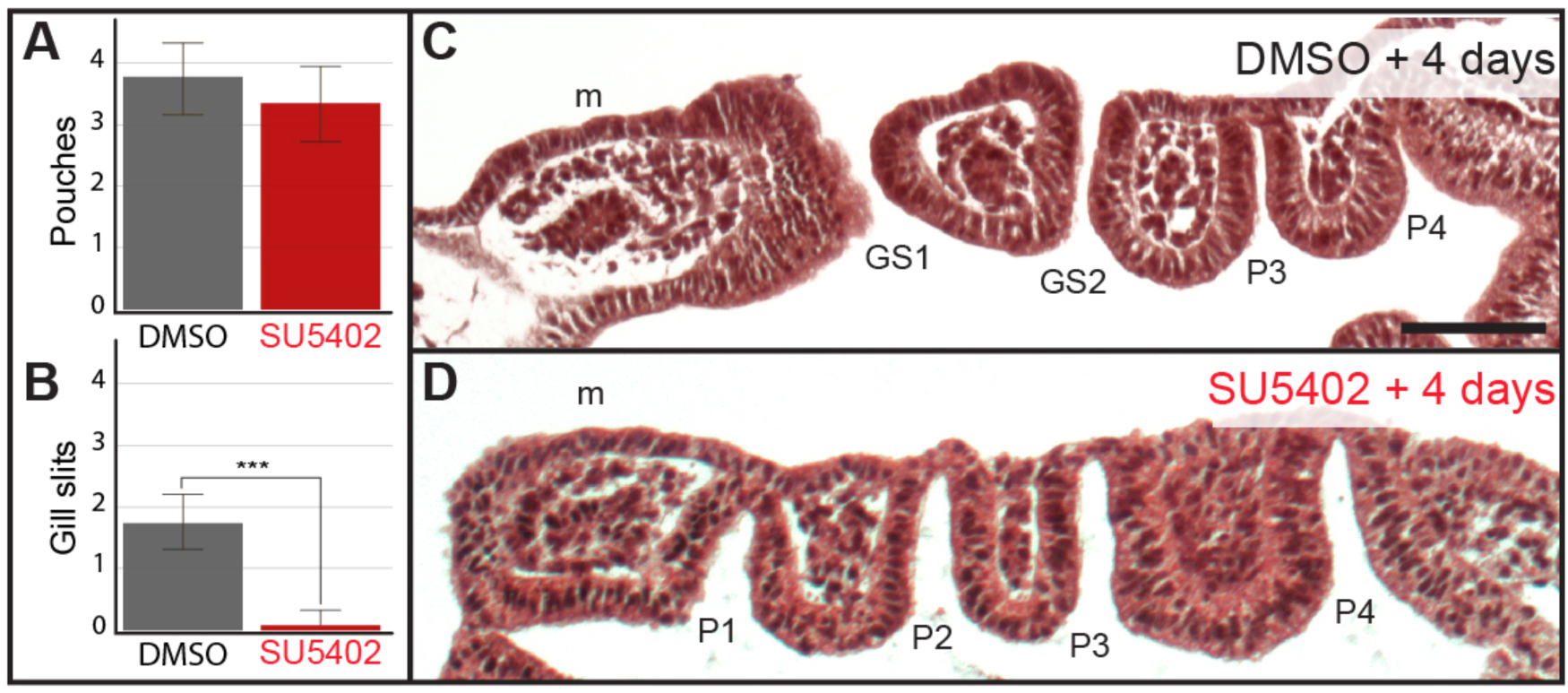
Fgf signalling is required for gill slit perforation in the skate. **(A)** There is no significant reduction in the number of pharyngeal pouches following four days of *in ovo* exposure to the Fgf receptor antagonist SU5402, relative to control. However, **(B)** there is a significant reduction in the number of gill slits following four days of *in ovo* exposure to SU5402, relative to control. **(C)** Representative histological section through an embryo after four days of exposure to DMSO. This embryo possesses two gill slits and two additional pharyngeal pouches that have not yet formed gill slits. **(D)** Representative histological section through an embryo after four days of exposure to SU5402. This embryo possesses four pharyngeal pouches, but no gill slits. *GS1-GS2*, gill slits 1-2; *P1–P4*, pharyngeal pouches 1– 4; *m*, mandibular arch. Scale bar = 100µm.

To determine the nature of the epithelial defects resulting from SU5402 treatment, we compared epithelial morphology between DMSO- and SU5402-treated embryos. In a representative control embryo, the 1^st^ (Fig. 7A) and 2^nd^ (Fig. 7B) gill slits had formed, while the 3^rd^ pharyngeal pouch was intercalated with the surface ectoderm (and the intercalated epithelium was beginning to undergo remodelling for perforation – Fig. 7C). In contrast, in a representative SU5402-treated embryo, the 1^st^ (Fig. 7D), 2^nd^ (Fig. 7E) and 3^rd^ pharyngeal pouches (Fig. 7F) had all intercalated with surface ectoderm, but this intercalated epithelium failed to undergo remodelling and gill slit perforation. With laminin and e-cadherin immunofluorescence (n=3 embryos), we observed that endodermal and ectodermal basement membrane degradation and epithelial cytoskeletal protrusions from the surface ectoderm toward pouch endoderm still occurred with SU5402 treatment (Fig. 8A,B), as did endoderm–ectoderm intercalation and re-establishment of basement membrane integrity within pharyngeal arches adjacent to the site of intercalation (Fig. 8C). These findings indicate that skate pharyngeal endoderm can still contact and intercalate with the ectoderm in the absence of FGF signalling, but that FGF signalling is required for final epithelial perforation and gill slit formation.

**Figure 7.**
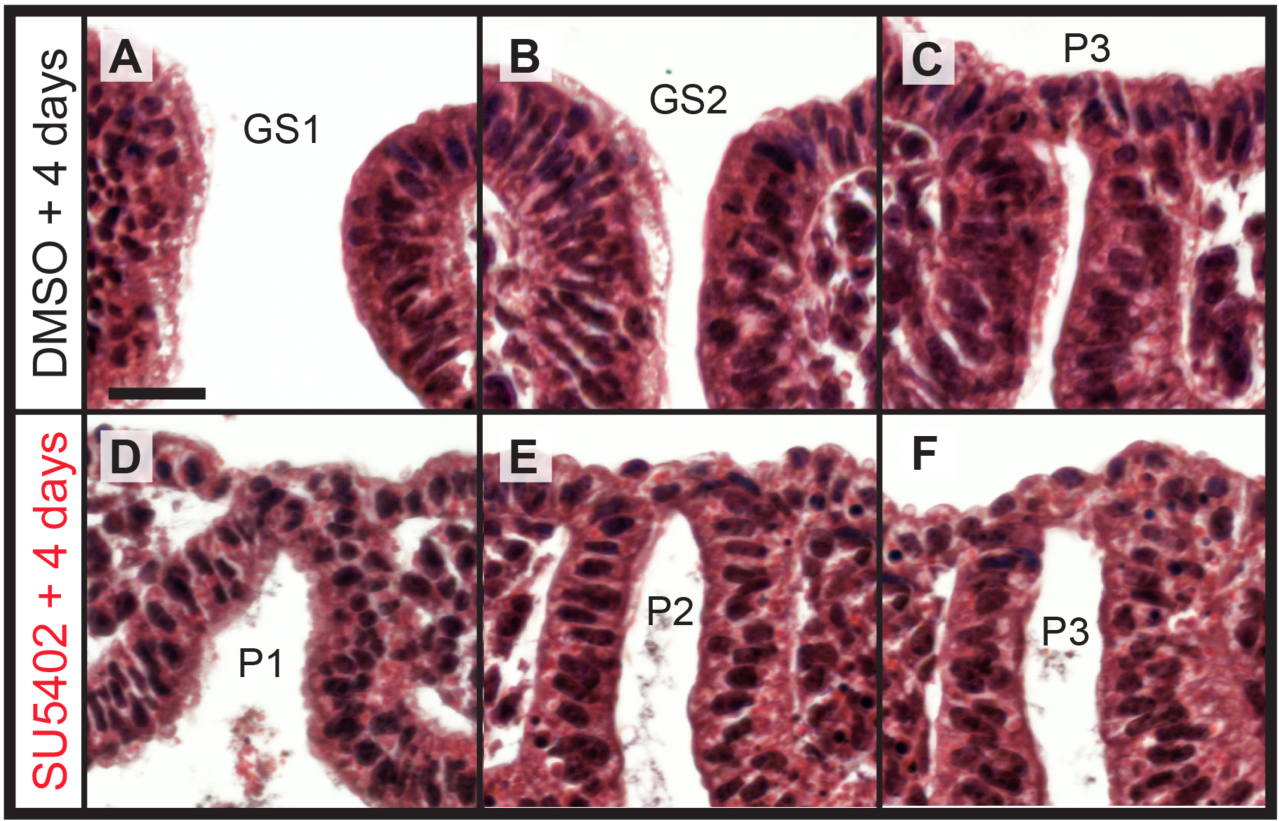
Fgf signalling is dispensable for epithelial intercalation but is required for gill slit perforation in the skate. In a representative skate embryo treated with DMSO *in ovo* for 4 days from S19/20, **(A)** gill slit 1 and **(B)** gill slit 2 have perforated, and **(C)** pouch 3 has intercalated with the surface ectoderm and perforation is initiating (*). By contrast, in a representative skate embryo treated with SU5402 *in ovo* for 4 days from S19/20, **(D)** pouch 1, **(E)** pouch 2 and **(F)** pouch 3 have all intercalated with the surface ectoderm, but gill slits have perforated. *GS1-GS2*, gill slits 1-2; *P1–P3*: pharyngeal pouches 1–3. All scale bar = 30µm.

**Figure 8.**
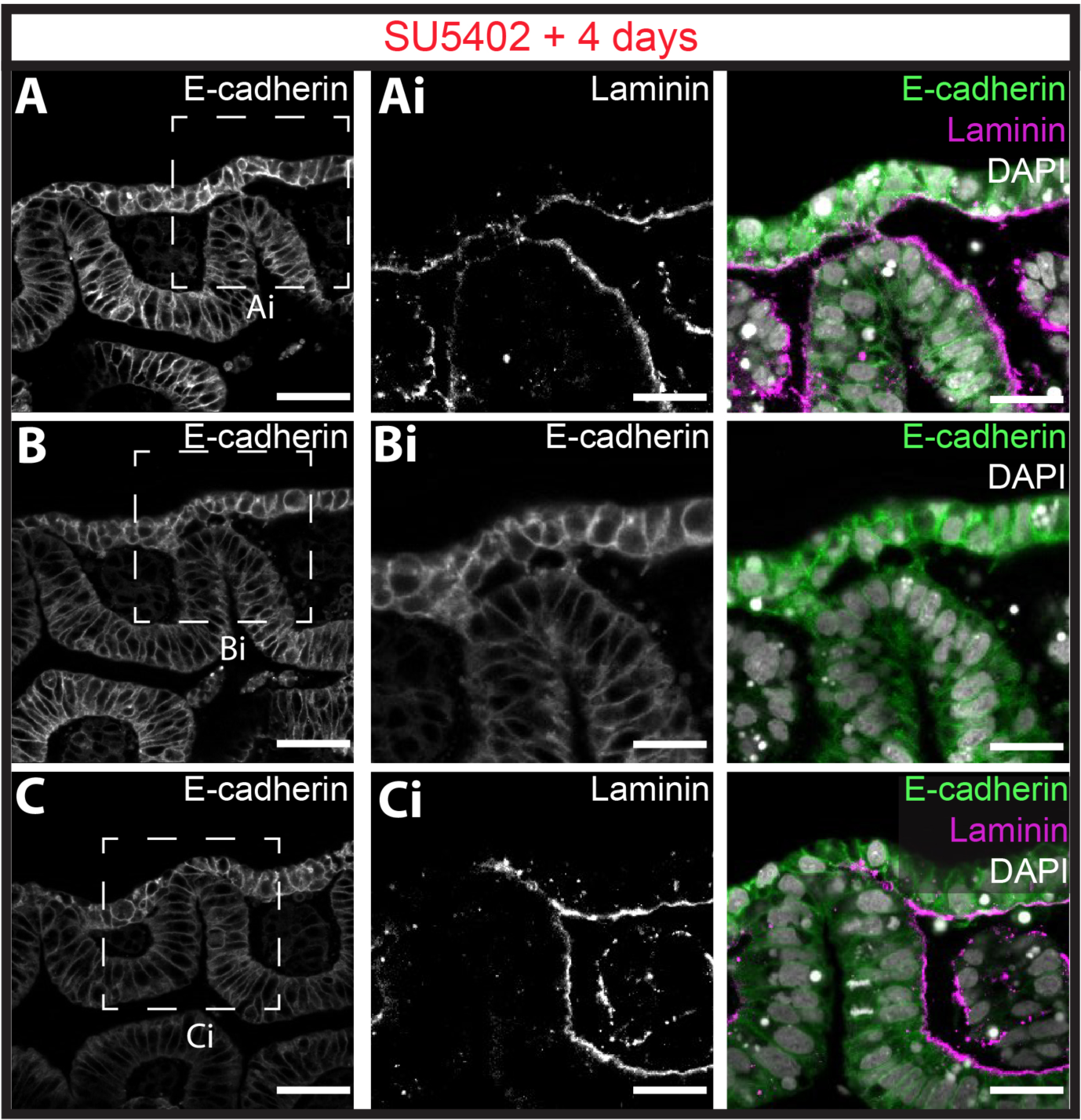
Fgf signalling dispensable for basement membrane degradation and endoderm-ectoderm intercalation but is required for gill slit perforation in the skate. Immunofluorescence for E-cadherin and laminin reveals that with SU5402 treatment **(A)** ectodermal basement membrane degradation, **(B)** ectodermal cytoskeletal protrusions toward the endoderm, **(C)** endoderm basement membrane degradation endoderm-ectoderm intercalation all occur normally. However, Fgf signalling is required for remodelling of intercalated epithelium and gill slit perforation. All pouches were imaged for n=3 SU5402-treated embryos, and representative images are shown here. Scale bars: A,B,C = 50µm; Ai,Bi,Ci = 25µm.

## Discussion

### Epithelial morphogenesis, perforation and gill slit formation

The endodermal–ectodermal interactions that we have described leading up to gill slit formation in skate differ from those previously reported from shark embryos in some key respects. Shone and Graham (2014) characterised epithelial morphology and basement membrane integrity during pharyngeal pouch morphogenesis in the small spotted catshark (*Scyliorhinus canicula*) and chick. They reported that both taxa exhibit disruption of basement membranes as pouch endoderm displaces and fuses with surface ectoderm. In the skate, we find that basement membrane degradation occurs first in the ectoderm, prior to endoderm–ectoderm contact. This is followed by ectodermal cytoskeletal protrusions that extend toward the endoderm and then ectoderm–endoderm contact. Similar degradation of the basement membrane with extension of epithelial protrusions leading to fusion of apposing epithelia has been previously described in other systems, such as the developing chick lung (Palmer and Nelson 2020) and may represent a general mechanism of epithelial fusion in vertebrates. Once the endodermal basement membrane has degraded, cells of the endoderm and ectoderm intercalate into a single epithelium, at which point it is no longer possible to distinguish distinct endodermal and ectodermal epithelia. Finally, we find that skate gill slit perforation occurs by remodelling and perforation of this intercalated epithelium, rather than by fusion with displaced ectoderm on either side of a persistent endodermal pouch.

In chick, perforation of the 2^nd^ pharyngeal pouch has been reported to occur by reorganisation of endodermal and ectodermal epithelia without cell death (Waterman, 1985) or via focal apoptosis at sites of endoderm–ectoderm contact (Shone and Graham, 2014). Our histological analysis of gill slit development and findings of negative TUNEL and cleaved caspase staining at points of endoderm–ectoderm contact indicates that, in skate, apoptosis does not contribute to endoderm–ectoderm fusion or gill slit perforation. Cell death during perforation of chick pharyngeal pouches was identified based on focal retention of Lysotracker Red at points of endoderm– ectoderm contact (Shone and Graham, 2014). It is possible that the Lysotracker Red staining is indicative of lysosomal activity related to basement membrane degradation rather than apoptosis. We therefore propose that epithelial remodelling, as described in skate and chick (Waterman, 1985) is the predominant and general process underlying epithelial perforation and gill slit formation in jawed vertebrates.

The formation of the vertebrate mouth offers another example of endoderm–ectoderm fusion and epithelial perforation during pharyngeal development (Chen et al., 2017). In *Xenopus*, mouth formation occurs at the juxtaposition of outer oral ectoderm and inner pharyngeal endoderm at the extreme anterior domain (EAD) of the embryonic head. This tissue interaction is followed by the breakdown of basement membranes, ectodermal cell death and intercalation of ectoderm and endoderm into a single epithelial layer that thins and perforates (Dickinson and Sive, 2006; 2009). Mouth development in *Xenopus* involves reciprocal signaling between the epithelium and adjacent neural crest-derived mesenchyme, with Wnt signals from the neural crest inducing convergent extension in overlying epithelium (Jacox et al., 2016). *Fgf8* is also expressed in the EAD (Chen et al., 2017), but its function in that context is not known. Broad similarities between processes of mouth and gill slit formation could reflect a shared underlying (i.e., serially homologous) program that directs interaction and fusion of endodermal and ectodermal epithelia in the vertebrate pharynx.

### Fgf signalling and vertebrate pharyngeal pouch development

Although expression of Fgf signalling components is a broadly conserved feature of pharyngeal arch patterning across vertebrates, the tissue specificity of these expression patterns during pharyngeal pouch development appears to vary across taxa. The invertebrate chordate amphioxus develops simple pharyngeal arches via endodermal pouches that are homologous with those of vertebrates, and these pouches show complex expression patterns of *Fgf8/17/18, Fgf9/16/20, FgfA* and *FgfC* (Bertrand et al., 2011). In the lamprey, *Fgf3* and *Fgf8* are expressed in the posterior region of each pharyngeal endodermal pouch with *Fgf8* additionally expressed in overlying pharyngeal ectoderm (Jandzik et al., 2014), and this regionalised expression of *Fgf8* and/or *Fgf3* within pharyngeal endormal pouches is conserved in chick (Veitch et al., 1999) and skate (reported here). In mouse *Fgf8* is expressed broadly throughout pharyngeal pouch endoderm and in overlying ectoderm, as well as in splanchnic mesoderm (Abu-Issa et al., 2002; Zbasnik et al., 2023), while in zebrafish *fgf8* is expressed in pharyngeal mesoderm during pouch formation (Choe and Crump, 2014). Taken together, these observations point to endodermal/ectodermally-derived Fgf signalling as an ancestral feature of developing chordate pharyngeal pouches, with additional and distinct mesodermal expression of *Fgf8* arising within the developing pharyngeal region of bony vertebrates.

Similarly variable are the functions of Fgf signalling during pharyngeal pouch development in vertebrates. In zebrafish, Fgf signalling from pharyngeal mesoderm has a well-established early function in guiding the lateral migration of presumptive pouch endoderm, and loss of Fgf signalling results in a failure of pharyngeal pouch formation (Crump et al., 2004; Choe and Crump, 2014). A loss of pharyngeal pouches also occurs with Fgf signalling inhibition in lamprey (Jandzik et al., 2014). We were unable to inhibit Fgf signalling prior to the onset of pouch formation with our *in ovo* pharmacological strategy, and this, combined with the relatively short duration of treatment, precluded us from rigorously testing for a role for Fgf signalling in early pharyngeal pouch development in skate. If, as in zebrafish, Fgf signalling functions as a chemoattractant for lateral migration of endoderm in the skate, it seems unlikely that this role is fulfilled by Fgf8 signalling from surface ectoderm (given the onset of ectodermal expression once pouch endoderm has already contacted surface ectoderm). Rather, this chemoattractant function could be fulfilled by other Fgf ligands expressed in pharynegal mesoderm and/or ectoderm. A broader survey of *Fgf* gene expression and a means of conducting earlier inhibition of Fgf signalling in skate embryos are needed to resolve this.

We do, however, present evidence of a previously undescribed later role for Fgf signalling in the remodelling and perforation of intercalated pharyngeal endoderm and ectoderm during gill slit formation in skate. During intercalation of pouch endoderm with surface ectoderm, we observe a sharp expression boundary between the *Fgf8*-/*Fgf3*-negative anterior and *Fgf8*-/Fgf3-positive posterior pouch endoderm. This boundary appears to correspond precisely with the future site of gill slit perforation and is coincident with the onset of *Fgf8* expression in overlying surface ectoderm. We therefore hypothesise that Fgf signalling is involved in determining the location of epithelial perforation prior to gill slit formation. Whether Fgf signalling within the endoderm or signalling from the surface ectoderm establishes the site of perforation is unclear, and further experimental work on tissue-specific roles of Fgf signalling during pharyngeal endodermal pouch and gill slit formation in cartilaginous and bony fish model systems will be needed to resolve this. This later role for Fgf signalling in pharyngeal pouch epithelial reorganisation and morphogenesis parallels, in some ways, the later (i.e., post-pouch formation) role for Fgf signalling in regulating pharyngeal pouch morphogenesis in mouse, and highlights the likely complex, varied, and multiphasic roles for Fgf signaling in governing epithelial behaviour during the establishment, shaping and remodelling of vertebrate pharyngeal arches and gill slits.

## Acknowledgements

The authors were funded by a Wellcome PhD studentship (214953/Z/18/Z) to JMR, by a Royal Society University Research Fellowship (UF130182 and URF\R\191007) and Royal Society Research Grant (RG140377) to JAG, and by Marine Biological Laboratory Institutional Funds to JAG.

---

**Supplemental Figure 1:**
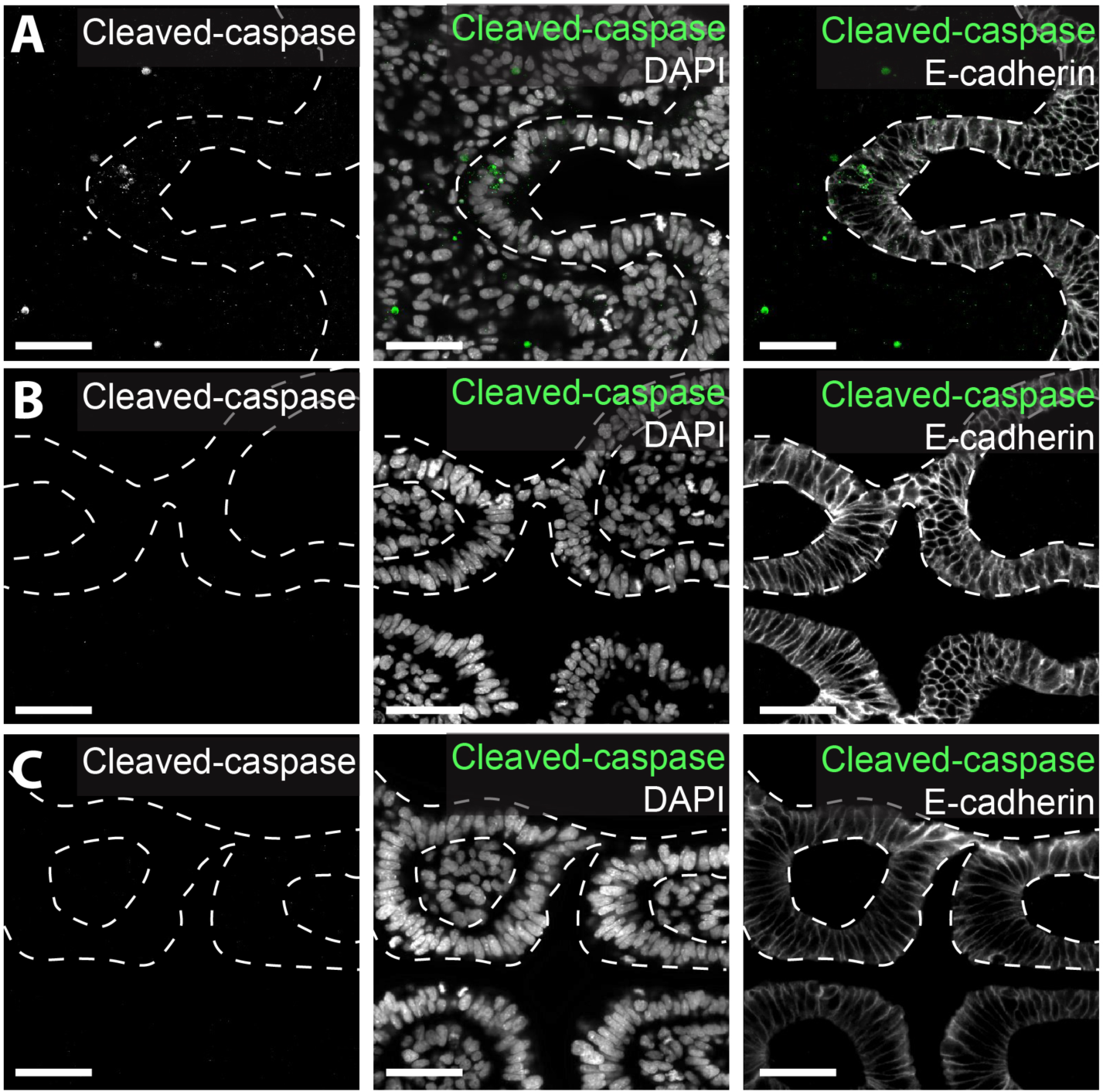
No cleaved caspase-3 expression during gill slit formation in the skate. Immunofluorescence for cleaved caspase-3 shows **(A)** a domain of positive staining in the rostral pharyngeal epithelium (consistent with our observation of TUNEL staining in this region) but **(B)** no cleaved caspase-3 expression in newly intercalated epithelium at the site of endoderm-ectoderm contact or **(C)** during gill slit perforation. All pouches were imaged in n=3 S22 embryos. All scale bars = 50µm.

**Supplemental Figure 2:**
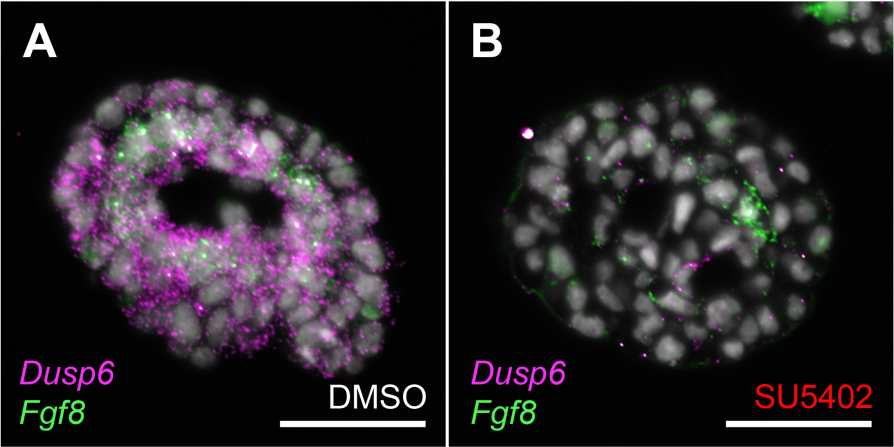
SU5402 treatment reduces expression of *Dusp6*, a transcriptional readout of Fgf signalling in the skate. Expression of *Dusp6* offers a strong transcriptional readout of Fgf signalling in the developing gill filaments of S24 skate embryos. To validate the effectiveness of our Fgf signalling inhibition strategy, we compared expression of *dusp6* in gill filaments using mRNA ISH by chain reaction (HCR) 24 hours after in ovo injection of **(A)** DMSO (control) or **(B)** SU5402. We observed a significant reduction in expression of *Dusp6* in gill filaments with SU5402 treatment. All scale bars = 50µm.

**Supplemental Figure 3:**
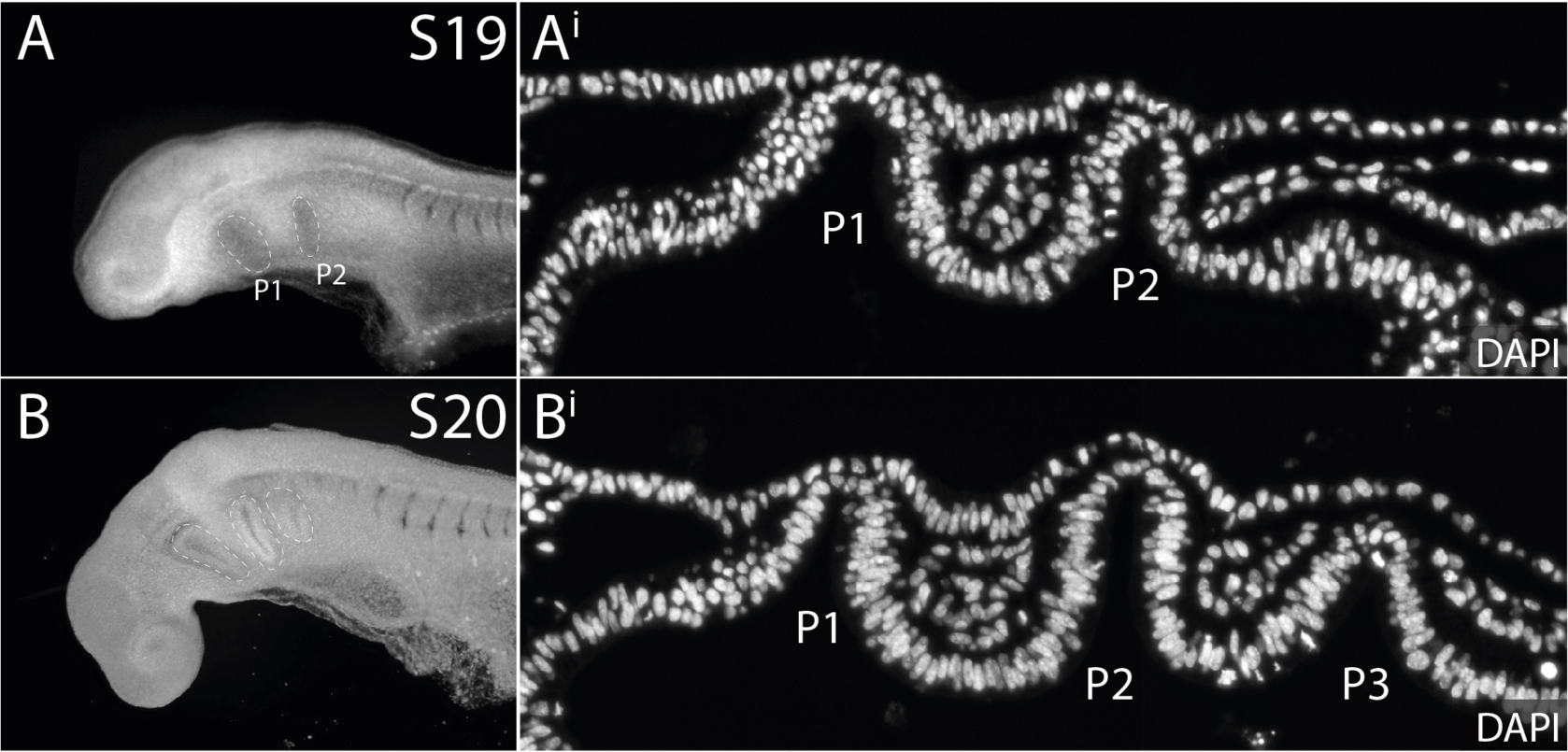
Pharyngeal pouch number in S19/20 skate embryos. Skate embryos were treated with SU5402 *in ovo* from S19/20. **(A)** At S19, skate embryos possess **(A^i^)** two pharyngeal pouches. **(B)** At S20, skate embryos possess **(B^i^)** three pharyngeal pouches. *P1–P3*: pharyngeal pouches 1–3.

## Notes

### Competing Interest Statement

The authors have declared no competing interest.

## References

Abu-Issa, R., Smyth, G., Smoak, I., Yamamura, K-I. & Meyers, E. N. 2002. Fgf8 is required for pharyngeal arch and cardiovascular development in the mouse. Development, 129, 4613–4625.

Ballard, W. W., Mellinger, J. & Lechenault, H. 1993. A series of normal stages for development of Scyliorhinus canicula, the lesser spotted dogfish (Chondrichthyes: Scyliorhinidae). Journal of Experimental Zoology, 267, 318–336.

Bertrand, S., Camasses, A., Somorjai, I., Belgacem, M. R., Chabrol, O., Escande, M. L., Pontarotti, P. & Escriva, H. 2011. Amphioxus FGF signaling predicts the acquisition of vertebrate morphological traits. Proc Natl Acad Sci U S A, 108, 9160–5.

Chen, J., Jacox, L. A., Saldanha, F. & Sive, H. 2017. Mouth development. Wiley Interdiscip Rev Dev Biol, 6.

Choe, C. P., Collazo, A., Trinh Le, A., Pan, L., Moens, C. B. & Crump, J. G. 2013. Wnt-dependent epithelial transitions drive pharyngeal pouch formation. Dev Cell, 24, 296–309.

Choe, C. P. & Crump, J. G. 2014. Tbx1 controls the morphogenesis of pharyngeal pouch epithelia through mesodermal Wnt11r and Fgf8a. Development, 141, 3583–93.

Choi, H. M. T., Schwarzkopf, M., Fornace, M. E., Acharya, A., Artavanis, G., Stegmaier, J., Cunha, A. & Pierce, N. A. 2018. Third-generation in situ hybridization chain reaction: multiplexed, quantitative, sensitive, versatile, robust. Development, 145.

Criswell, K. E. & Gillis, J. A. 2020. Resegmentation is an ancestral feature of the gnathostome vertebral skeleton. Elife, 9.

Crump, J. G., Maves, L., Lawson, N. D., Weinstein, B. M. & Kimmel, C. B. 2004. An essential role for Fgfs in endodermal pouch formation influences later craniofacial skeletal patterning. Development, 131, 5703–16.

Dickinson, A. J. & Sive, H. 2006. Development of the primary mouth in Xenopus laevis. Dev Biol, 295, 700–13.

Dickinson, A. J. & Sive, H. L. 2009. The Wnt antagonists Frzb-1 and Crescent locally regulate basement membrane dissolution in the developing primary mouth. Development, 136, 1071–81.

Edlund, R. K., Ohyama, T., Kantarci, H., Riley, B. B. & Groves, A. K. 2014. Foxi transcription factors promote pharyngeal arch development by regulating formation of FGF signaling centers. Developmental Biology, 390, 1–13.

Gillis, J. A., Modrell, M. S., Northcutt, R. G., Catania, K. C., Luer, C. A. & Baker, C. V. 2012. Electrosensory ampullary organs are derived from lateral line placodes in cartilaginous fishes. Development, 139, 3142–6.

Gillis, J. A. & Tidswell, O. R. 2017. The Origin of Vertebrate Gills. Curr Biol, 27, 729–732.

Graham, A., Poopalasundaram, S., Shone, V. & Kiecker, C. 2019. A reappraisal and revision of the numbering of the pharyngeal arches. J Anat, 235, 1019–1023.

Graham, A. & Smith, A. 2001. Patterning the pharyngeal arches. Bioessays, 23, 54–61.

Grevellec, A. & Tucker, A. S. 2010. The pharyngeal pouches and clefts: Development, evolution, structure and derivatives. Semin Cell Dev Biol, 21, 325–32.

Houssin, N. S., Bharathan, N. K., Turner, S. D. & Dickinson, A. J. 2017. Role of JNK during buccopharyngeal membrane perforation, the last step of embryonic mouth formation. Dev Dyn, 246, 100–115.

Jacox, L., Chen, J., Rothman, A., Lathrop-Marshall, H. & Sive, H. 2016. Formation of a “Pre-mouth Array” from the Extreme Anterior Domain Is Directed by Neural Crest and Wnt/PCP Signaling. Cell Rep, 16, 1445–1455.

Jandzik, D., Hawkins, M. B., Cattell, M. V., Cerny, R., Square, T. A. & Medeiros, D. M. 2014. Roles for FGF in lamprey pharyngeal pouch formation and skeletogenesis highlight ancestral functions in the vertebrate head. Development, 141, 629–38.

Kopinke, D., Sasine, J., Swift, J., Stephens, W. Z. & Piotrowski, T. 2006. Retinoic acid is required for endodermal pouch morphogenesis and not for pharyngeal endoderm specification. Dev Dyn, 235, 2695–709.

Loo, D. T. 2011. In situ detection of apoptosis by the TUNEL assay: an overview of techniques. Methods Mol Biol, 682, 3–13.

Mohammadi, M., Mcmahon, G., Sun, L., Tang, C., Hirth, P., Yeh, B. K., Hubbard, S. R. & Schlessinger, J. 1997. Structures of the tyrosine kinase domain of fibroblast growth factor receptor in complex with inhibitors. Science, 276, 955–60.

Mole, M. A., Galea, G. L., Rolo, A., Weberling, A., Nychyk, O., De Castro, S. C., Savery, D., Fassler, R., Ybot-Gonzalez, P., Greene, N. D. E. & Copp, A. J. 2020. Integrin-Mediated Focal Anchorage Drives Epithelial Zippering during Mouse Neural Tube Closure. Dev Cell, 52, 321–334 e6.

Nishimura, T., Honda, H. & Takeichi, M. 2012. Planar cell polarity links axes of spatial dynamics in neural-tube closure. Cell, 149, 1084–97.

Noden, D. M. 1983. The role of the neural crest in patterning of avian cranial skeletal, connective, and muscle tissues. Dev Biol, 96, 144–65.

O’neill, P., Mccole, R. B. & Baker, C. V. 2007. A molecular analysis of neurogenic placode and cranial sensory ganglion development in the shark, Scyliorhinus canicula. Dev Biol, 304, 156–81.

Palmer, M. A., Nelson, C. M. 2018. Epithelial tube fusion as a mechanism for the development of complex lumen-containing organs. Trends in Developmental Biology.

Palmer, M. A., Nelson, C. M. 2020. Fusion of airways during avian lung development constitutes a novel mechanism for the formation of continuous lumena in multicellular epithelia. Developmental Dynamics, 249: 1318–1333.

Piotrowski, T., Ahn, D. G., Schilling, T. F., Nair, S., Ruvinsky, I., Geisler, R., Rauch, G. J., Haffter, P., Zon, L. I., Zhou, Y., Foott, H., Dawid, I. B. & Ho, R. K. 2003. The zebrafish van gogh mutation disrupts tbx1, which is involved in the DiGeorge deletion syndrome in humans. Development, 130, 5043–52.

Piotrowski, T. & Nusslein-Volhard, C. 2000. The endoderm plays an important role in patterning the segmented pharyngeal region in zebrafish (Danio rerio). Dev Biol, 225, 339–56.

Rolo, A., Savery, D., Escuin, S., De Castro, S. C., Armer, H. E., Munro, P. M., Mole, M. A., Greene, N. D. & Copp, A. J. 2016. Regulation of cell protrusions by small GTPases during fusion of the neural folds. Elife, 5, e13273.

Sabapathy, K., Jochum, W., Hochedlinger, K., Chang, L., Karin, M. & Wagner, E. F. 1999. Defective neural tube morphogenesis and altered apoptosis in the absence of both JNK1 and JNK2. Mech Dev, 89, 115–24.

Sai, X. & Ladher, R. K. 2008. FGF signaling regulates cytoskeletal remodeling during epithelial morphogenesis. Curr Biol, 18, 976–81.

Shone, V. & Graham, A. 2014. Endodermal/ectodermal interfaces during pharyngeal segmentation in vertebrates. J Anat, 225, 479–91.

Sleight, V. A. & Gillis, J. A. 2020. Embryonic origin and serial homology of gill arches and paired fins in the skate, Leucoraja erinacea. Elife, 9, 2020.07.02.183665.

Veitch, E., Begbie, J., Schilling, T. F., Smith, M. M. & Graham, A. 1999. Pharyngeal arch patterning in the absence of neural crest. Curr Biol, 9, 1481–4.

Waterman, R. E. 1985. Formation and perforation of closing plates in the chick embryo. Anat Rec, 211, 450–7.

Xie, Y., Su, N., Yang, J., Tan, Q., Huang, S., Jin, M., Ni, Z., Zhang, B., Zhang, D., Luo, F., Chen, H., Sun, X., Feng, J. Q., Qi, H. & Chen, L. 2020. FGF/FGFR signaling in health and disease. Signal Transduct Target Ther, 5, 181.

Zbasnik, N. & Fish, J.L. 2023. *Fgf8* regulates first pharyngeal arch segmentation through pouch-cleft interactions. Front Cell Dev Biol. Doi. 10.3389/fcell.2023.1186526.

